# Dose response relationships for gamma radiation induced chromosomal instability

**DOI:** 10.1101/224741

**Authors:** Y.A. Eidelman, S.V. Slanina, V.S. Pyatenko, I.K. Khvostunov, S.G. Andreev

## Abstract

Different cell lines demonstrate various dose response for radiation-induced chromosomal instability (RICI). To clarify the origin of differences we analyzed own and published data on RICI for four cell lines, V79, TK6, WTK1 and CHO-K1 on the basis of the mechanistic RICI model. We conclude that observable dose-response shapes, both plateau-like and strong dose dependent behavior, may be jointly explained by the same model of RICI. Mechanistic modeling reveals that a variation of certain set of RICI parameters leads to strong modification of dose-response curve.

## INTRODUCTION

It is widely believed that radiation induced chromosomal instability (RICI) is dose-independent [1]. In particular, chromosomal aberrations like delayed dicentrics showed dose independence [1, 2]. The goal of the study was to examine the generality of this phenomenon and its sources. For this purpose we collated the most reliable data on delayed dicentrics by using the following criteria: sufficient statistics, numerous dose points and availability of dose-dependence along with time-dependence. The whole data pool including own experiments was analyzed by means of computer model of RICI developed [3, 4].

## METHODS

The *in vitro* experimental study and computer simulations were carried out as described elsewhere [3–5].

## RESULTS

Fig. 1 shows the collection of experimental data on dose-response curves for delayed dicentrics in three different cell lines. The data indicate a dose-independent (plateau-like) behavior at intermediate and high doses for V79 and TK6 cells, and at all doses for WTK1 cells.

**Fig.1.**
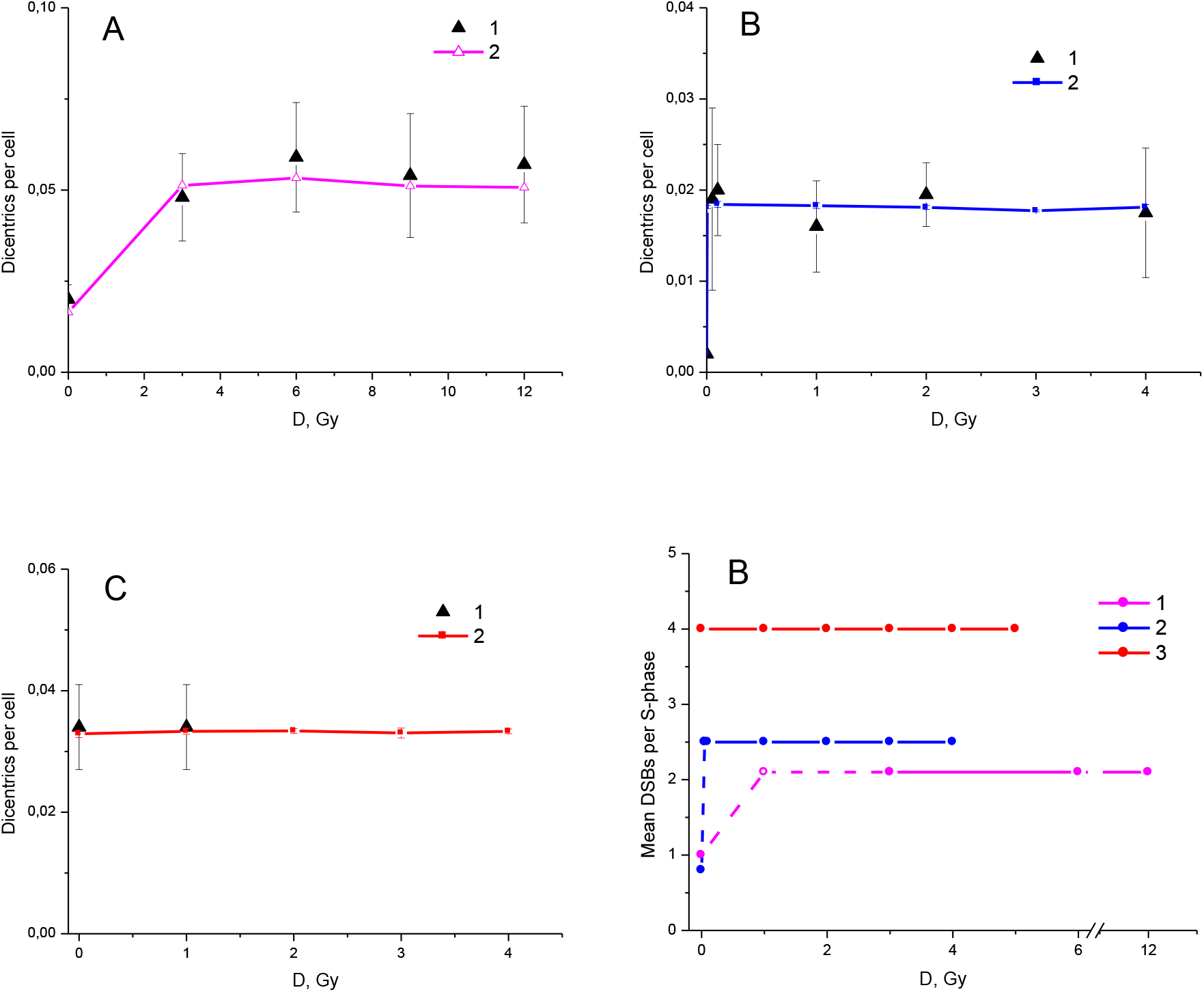
Experimental and simulated frequencies of dicentrics at late harvest times as a function of dose. A: V79 cells. The frequency of dicentrics was measured on the 14th day after irradiation, the cells were reseeded on the 7th and 13th days. 1 - experiment [6]. Error bars are standard deviations. 2 – RICI modeling. B: TK6 cells. Measurement on a long time (~ 35 generations). 1 - experiment [7]. Error bars are standard errors. The cell density was maintained at 2-10x10^5^ cells / ml, which approximately corresponds to a two-day reseeding schedule. 2 - RICI modeling with reseeding every two days. C: WTK1 cells, experiment [7] and simulation. The measurement conditions and designations are the same as in B. D: theoretically reconstructed DSB generation rate in the progeny for different cell lines as a function of dose. Cell lines: 1 - V79, 2 - TK6, 3 - WTK1.

The computer simulation using RICI model was carried out for the selected three cell lines for which the data are available and of sufficient quality. The simulation results agreed with the experimental points well (Fig.1). This fitting allowed to estimate the generation rate of persistent DSBs per S phase in the progeny of irradiated cells which is one of the main RICI model parameters. The rate proved to be dose-independent in the range of intermediate and high doses for V79 and TK6 cells. For WTK1 cells, DSB generation rate is dose-independent at whole dose range, including zero value.

The following questions arise from these findings: (1) what mechanism is responsible for the plateau on the dicentric dose-response curves? (2) how can the data be interpreted on the basis of the theoretical model of RICI? The naive explanation of RICI dose-independence is that it arises from dose-independence of DSBs generation rate and just repeats its shape. However, this is not the case, as can be seen from Fig.2.

**Fig.2.**
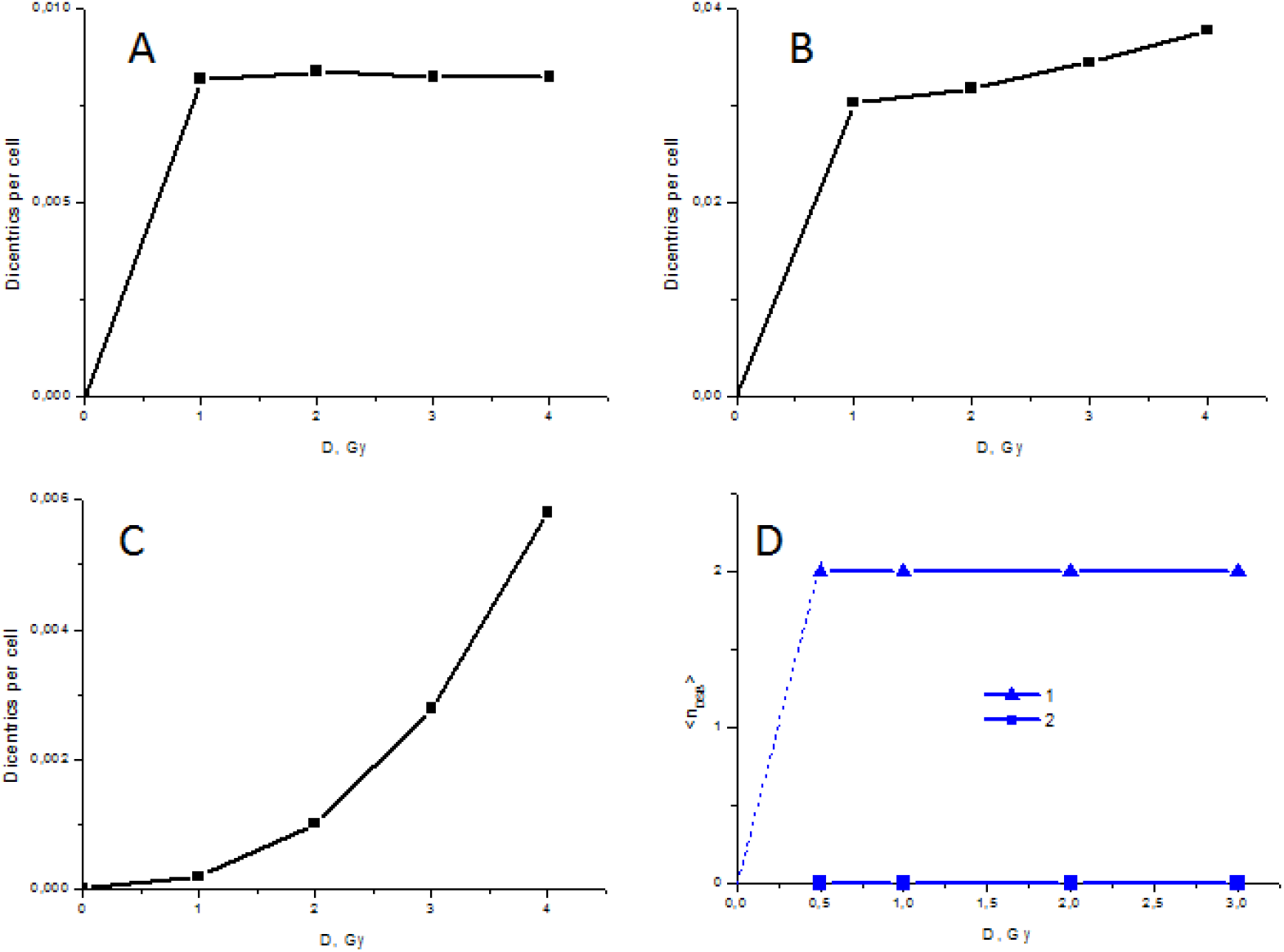
The frequency of delayed dicentrics, simulated by means of the RICI model, and corresponding generation rate of DSB as a function of dose using different parameter sets. Harvest time is 10 days post-irradiation. A: DSB generation 2 per S phase, probability of bridge breakage 0.4. B: DSB generation 2 per S phase, probability of bridge breakage 0.7. C: no DSB generation, probability of bridge breakage 0.7. D: dose dependencies of DSB generation rate in the progeny. 1 - corresponding to panels A and B. 2 - corresponding to panel C.

Panels A-C in Fig.2 reveal that the simulated dose-response for delayed dicentrics strongly depends on combination of mechanistic parameters, although DSB generation rate remains dose-independent.

The type of dose dependence becomes a plateau for the particular set of parameters that reflects certain relationships between rates of generation and transfer of chromosome lesions through the cell cycle in the progeny of irradiated cells. The non-zero control existing for all cell lines (Fig. 1) was not taken into account in Fig. 2.

To examine the conclusion drawn from Fig.1 of RICI dose-response curve independence we carried out the experimental study of RICI in Chinese hamster ovary cells CHO-K1. The experimental data on delayed dicentrics are shown in Fig. 3.

**Fig. 3.**
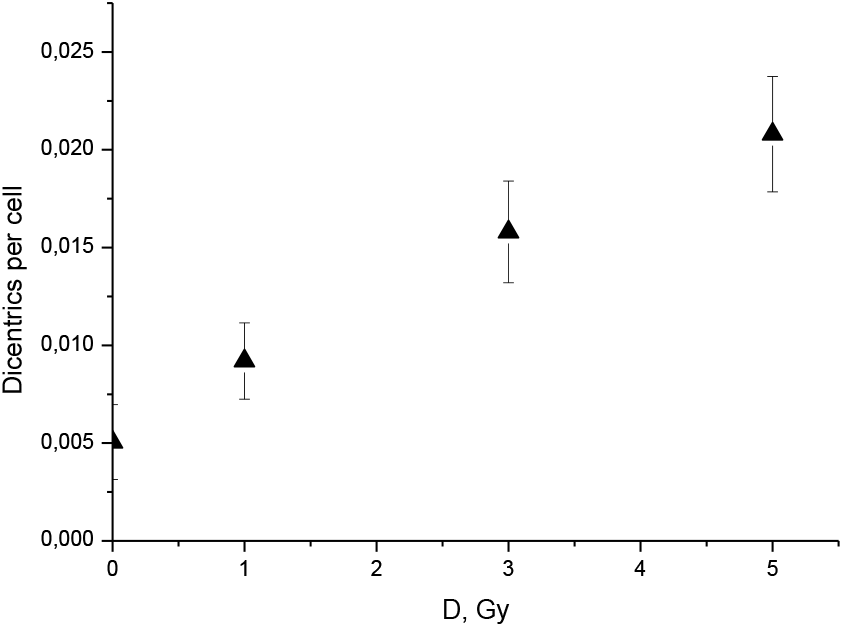
Dose dependence of delayed dicentrics in CHO-K1 cells, observed 17 days after gamma exposure.

The quasi-linear shape of the experimental dose-response curve shown in Fig. 3 differs markedly from the theoretical predictions shown in Fig 2. To determine the reason for this difference we performed several *in vitro* and *in silico* experiments, see Fig.4.

**Fig.4.**
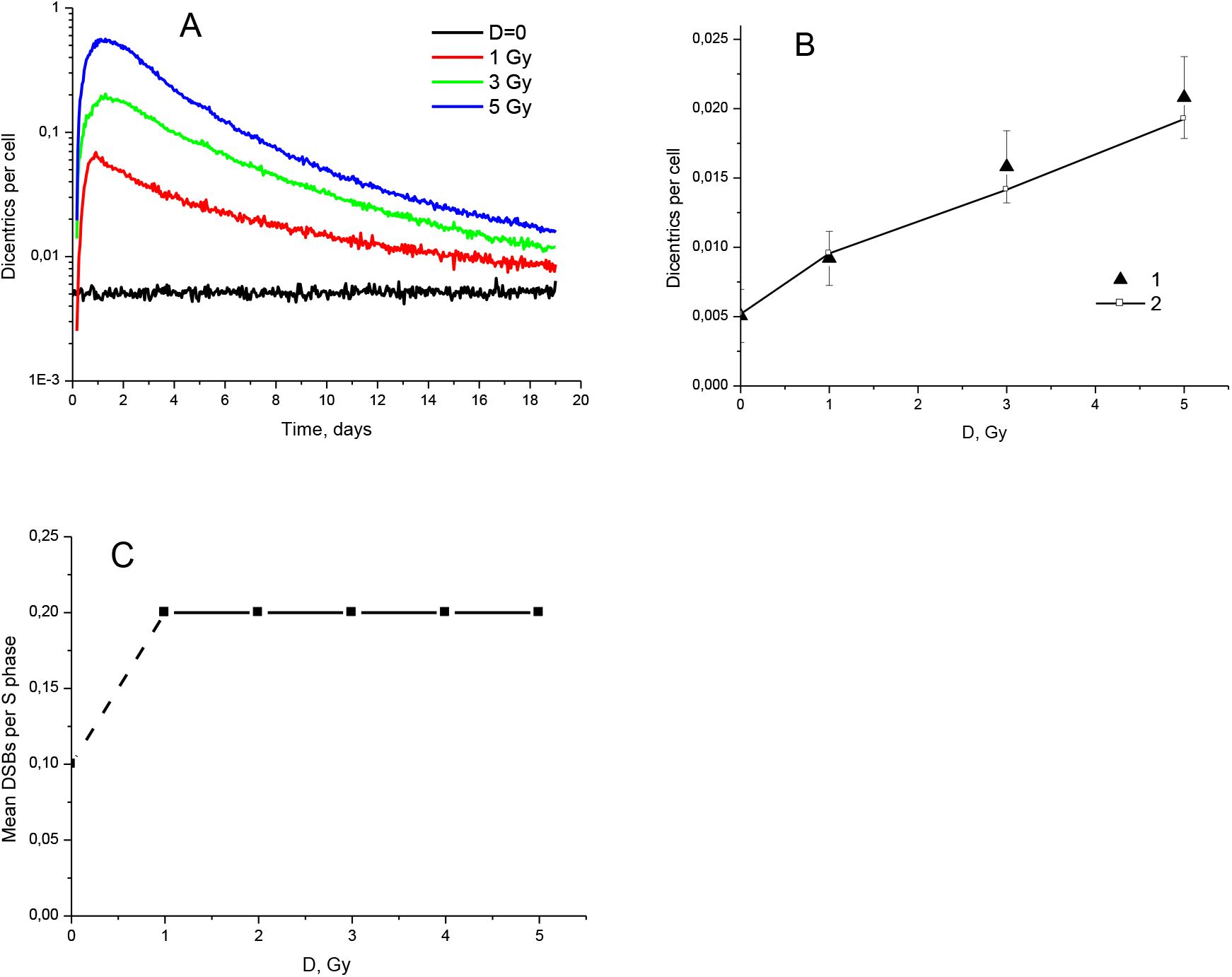
Dose-time dependencies of dicentric frequency in CHO-K1 cells. A: time dependencies for different doses, simulation by RICI model. B: Dose dependence at 17 days after exposure. 1 – experiment (same as in Fig.3), 2 – simulation by RICI model. C: theoretically reconstructed DSB generation rate in the progeny for different cell lines as a function of dose.

The analysis showed that for a certain form of dynamic curves (Fig. 4A), the quasilinear dose dependence was indeed obtained (Fig. 4B), in spite of the fact that DSB generation and interaction rates in the progeny do not depend on dose and time (Fig.4C).

Thus, the RICI model demonstrates the ability to describe both the dose plateau and the dose dependence observed in the experiment, Figs 1, 3, 4.

### Conclusions

In addition to the dose-independence of RICI at moderate and high doses, a new dose-response curve shape was experimentally observed in CHO-K1 cells, dose-increase in the frequency of dicentrics in the progeny of irradiated cells.

For the first time, a theoretical explanation of this dose dependence of RICI was presented, which was in concert with the available data on dicentric dynamics, as well as internally self-consistent. The model predicts conditions or range of mechanistic parameters in which the dose independence or dose dependence of the dicentric frequency may be observed.

## ACKNOWLEDGEMENTS

The present work was supported by Russian Foundation for Basic Research grant 14-01-00825 to S.A. S.A. acknowledges support from the MEPhI Academic Excellence Project (Contract No. 02.a03.21.0005).

## REFERENCES

1 United Nations Scientific Committee on the Effects of Atomic Radiation (UNSCEAR). Effects of ionizing radiation. UNSCEAR 2006 Report.

2 Morgan W.F., Sowa M.B. Non-targeted effects of ionizing radiation: implications for risk assessment and the radiation dose response profile. Health Phys. 2009, 97, 426–432.

3 Andreev S.G., Eidelman Y.A., Salnikov I.V., Slanina S.V. Modeling Study of Dose-Response Relationships for Radiation-Induced Chromosomal Instability. Dokl. Biochem. Biophys. 2013, 451, 171–175.

4 Eidelman Y.A., Slanina S.V., Pyatenko V.S., Andreev S.G. DNA Damage Induced Chromosomal Instability. Computational modeling view. BioRxiv preprint 2017, doi: http://dx.doi.org/10.1101/218826.

5 Pyatenko V.S., Eidelman Y.A., Khvostunov I.K., Andreev S.G. Radiation-induced chromosomal instability under constrained growth of irradiated cells. Dokl.Biochem.Biophys. 2013, 451, 190–193.

6 Jamali M., Trott K.R. Persistent increase in the rates of apoptosis and dicentric chromosomes in surviving V79 cells after X-irradiation. Int. J. Radiat. Biol. 1996, 70, 705–709.

7 Schwartz J.L., Jordan R., Evans H.H., Lenarczyk M., Liber H. The TP53 dependence of radiation-induced chromosome instability in human lymphoblastoid cells. Radiat. Res. 2003, 159, 730–736.

